# The intrinsic neonatal hippocampal network: rsfMRI findings

**DOI:** 10.1101/823500

**Authors:** Athena L. Howell, David E. Osher, Jin Li, Zeynep M. Saygin

**Affiliations:** The Ohio State University, 43210

## Abstract

Many adults cannot voluntarily recall memories before the ages of 3-5, a phenomenon referred to as “infantile amnesia” The development of the hippocampal network likely plays a significant part in the emergence of the ability to form long-lasting memories. In adults, the hippocampus has specialized and privileged connections with certain cortical networks, which presumably facilitate its involvement in memory encoding, consolidation, and retrieval. Is the hippocampus already specialized in these cortical connections at birth? And are the topographical principles of connectivity (e.g. long-axis specialization) present at birth? We analyzed resting-state hippocampal connectivity in neonates scanned within one week of birth (Developmental Human Connectome Project) and compared them to adults (Human Connectome Project). We explored the connections of the whole hippocampus and its long-axis specialization to seven canonical cortical networks. We found that the neonatal hippocampal networks show clear immaturity at birth: adults showed hippocampal connectivity that was unique for each cortical network, whereas neonates showed no differentiation in hippocampal connectivity across these networks. Further, neonates lacked long-axis specialization (i.e., along anterior-posterior axis) of the hippocampus in its differential connectivity patterns to the cortical networks. This immaturity in connectivity may contribute to immaturity in memory formation in the first years of life.

**“New and Noteworthy”:** While animal data, and anatomical and behavioral human data from young children suggest that the hippocampus is immature at birth, to date, there are no direct assessments of human hippocampal functional connectivity (FC) very early in life. Our study explores the FC of the hippocampus to the cortex at birth, allowing insight into the development of human memory systems.

## Introduction

Many adults cannot voluntarily recall memories before the ages of 3-5, a phenomenon referred to as “infantile amnesia” (Alberini & Travaglia, 2017). One potential reason for this is that the hippocampus (the primary brain structure responsible for episodic memory formation in adults) and its connections with the rest of the brain may be immature at birth. Indeed, the hippocampus does appear to be immature at birth; evidence in macaques suggests it continues to mature after one year of age (roughly age 3-5 in humans) (Jabés et al., 2011) and human data indicates that volumetric and structural changes in the hippocampus continue through childhood (DeMaster et al., 2014; Gilmore et al., 2012; Seress 2007). Further, episodic memory performance may be influenced by changes in the patterns of hippocampal connectivity from middle childhood to adulthood, including along the long-axis of the hippocampus (Blankenship et al., 2017; DeMaster et al., 2014; Ghetti et al., 2010; Gogtay et al., 2006; Poppenk & Moscovitch, 2011; Riggins et al., 2016). At younger ages, hippocampal gray matter volume has been linked to early language ability (Can et al., 2013) and one recent study showed potential hippocampal activation for learned items in 2-year old toddlers (Prabhakar et al., 2018). However, the intrinsic connectivity of hippocampus very early in life is less well understood. Therefore, an understanding of the hippocampal network at birth and its development may lead to greater understanding of memory development.

Recently, Wael and colleagues (2018) showed the hippocampus has a clear intrinsic pattern of functional connectivity (FC) to a set of cortical networks in adults. Specifically, they showed higher (i.e. most positive) connectivity from the hippocampus to the Default Mode and Limbic networks and lowest (i.e. least positive) connectivity to the Frontoparietal and Ventral Attention networks (from Yeo et al., 2011), Further, this connectivity pattern differed between the anterior and posterior portions of the hippocampus, with anterior hippocampus showing larger differences in connectivity to the networks than posterior hippocampus. This so-called long-axis specialization of the hippocampus is consistent with previous research showing that the anterior and posterior hippocampus display different patterns of structural and functional connectivity and may be uniquely activated in response to cognitive, memory and spatial demands (for reviews see Poppenk et al., 2013, Strange et al., 2014). The development of the hippocampal network and the long-axis gradient likely plays a significant part in the emergence of the ability to form long-lasting memories. For instance, the work of Riggins et al. (2016) examines the relationship of anterior/posterior connectivity and episodic memory in 4- and 6-year old children and finds developmental differences even between these two ages. Although this work in young children is notable, the fact remains that we know very little about the hippocampus, its connections, and its relationship to memory-formation during the earliest stages of life.

To this end, we compared the resting-state hippocampal connectivity patterns to a set of cortical networks in neonates and adults. Resting state connectivity, determined by spontaneously correlated activity of disparate brain regions, is used as a reliable marker of intrinsic functional connectivity (FC) between brain regions (Biswal et al., 1995; Raichle, 2009; Smith, 2013; Sporns, 2013); further, FC at rest is predictive of task-based activity (Cole et al., 2014; Osher et al., 2019; Smith et al., 2009; Tobyne et al., 2018).

More recently, developmental studies using FC have shown the FC of some networks is mature at birth while others take months or longer to become adultlike (for reviews see Gao et al., 2017 and Grayson & Fair, 2017). In particular, multiple studies indicate the connectivity of visual and somatomotor networks is not only functional but highly adult-like at birth (Gao et al., 2015b; Lin et al., 2008; Liu et al., 2008). Other areas, such as the default mode network, dorsal attention network, frontoparietal network, and some perceptual regions show relatively immature functional and structural characteristics at birth and experience large modifications postnatally (Gao et al., 2015b; Natu et al., 2019), although the frontoparietal network may have important functional roles even within the first year of life (e.g. Linke et al., 2018).

To assess hippocampal maturity at birth, we analyzed FC between seven intrinsic networks and the hippocampus as a whole as well as along the hippocampal long-axis in both neonates and adults. We also compared neonatal vs. adult hippocampal connectivity to the cortex at a finer, voxelwise scale. Based on previous literature suggesting the immaturity of the hippocampus at birth, we hypothesized that neonates would differ from adults in their hippocampal connectivity to the cortex, particularly to the more immature networks (e.g. default mode and frontoparietal).

## Materials and Methods

### Participants

#### Neonates

Neonatal data comes from the initial release of the Developing Human Connectome Project (dHCP) (http://www.developingconnectome.org, Markopoulos et al., 2018). Neonates were recruited and imaged in London at the Evelina Neonatal Imaging Centre after gathering informed parental consent to image and release the data. The study was approved by the UK Health Research Authority. 40 neonates were included in our analyses (15 female, 36-44 weeks old at scan).

#### Adults

Adult data comes from the Human Connectome Project (HCP), WU-Minn HCP 1200 Subject Data Release (https://www.humanconnectome.org/study/hcp-young-adult, Van Essen et al., 2013). Participants were scanned at Washington University in St. Louis (WashU). We included 40 participants in our analyses (15 female; 20-35 years old). These adult participants were motion-matched to the neonates. Specifically, we matched each neonatal participant with an adult from the HCP dataset with the same gender who showed the most similar motion parameter (i.e., framewise displacement, FD). While the resulting group of adults were motion-matched to the neonates, we found that the groups were significantly different in the average gray-matter tSNR, with neonates exhibiting higher tSNR values (*t(78)=*-6.8774, *p=*1.3469×10^−9^). To ensure that any results were not driven by tSNR differences between groups, we identified an additional group of HCP adults whose tSNR was matched to the tSNR of the neonates *(t(78*)=-1.5237, *p*=.132) and replicated our results (see Extended Data, Figure 1-1).

### MRI Acquisition

#### Neonates

All acquisition information comes from the dHCP data release documentation. Imaging was carried out on a 3T Philips Achieva (running modified R3.2.2 software) using an imaging system specifically designed for neonates with a 32 channel phased array head coil (Hughes, E.J., et al.). Neonates were scanned during natural sleep; resting-state FC patterns have been shown to stay largely consistent while awake, asleep, or under anesthesia (Liu et al., 2015; Larson-Prior et al., 2009).

#### Resting-state fMRI

High temporal resolution fMRI developed specifically for neonates was collected using multiband (MB) 9x accelerated echo=planar imaging (TE/TR=38/392ms, voxel size = 2.15 x 2.15 x 2.15mm^3^). The resting state scan lasted approximately 15 minutes and consisted of 2300 volumes for each run. No in-plane acceleration or partial Fourier transform was used. Single-band reference scans with bandwidth matched readout and additional spin-echo acquisitions were also acquired with both AP/PA fold-over encoded directions.

#### Anatomical MRI

High-resolution T2-weighted and inversion recovery T1-weighted multi-slice fast spin-echo images were acquired with in-plane resolution 0.8 x 0.8mm^2^ and 1.6mm slices overlapped by 0.8mm (T2-weighted: TE/TR= 156/12000ms; T1 weighted: TE/TR/TI = 8.7/4795/1740ms)

#### Adults

All acquisition information comes from the HCP data release documentation. Scanning for the 1200 WU-Minn HCP subject was carried out on a customized 3T Connectome Scanner adapted from a Siemens Skyra (Siemens AG, Erlanger, Germany), equipped with a 32-channel Siemens receiver head coil and a “body” transmission coil specifically designed by Siemens to accommodate the smaller space (due to special gradients) of the WU-Minn and MGH-UCLA Connectome scanners.

#### Resting-State fMRI

Participants were scanned using the Gradient-echo EPI sequence (TE/TR = 33.1/720ms, flip angle = 52°, 72 slices, voxel size = 2 x 2 x 2mm^3^). Scanning lasted approximately 15 minutes consisting of 1200 volumes for each run. Each participant finished two resting-state fMRI sessions. For each session, two phases were encoded: one right-to-left (RL) and the other left-to-right (LR). For our analyses, we used the LR phase encoding from the first session. Participants were instructed to relax and keep their eyes open and fixated on a bright, projected cross-hair against a dark background.

#### Anatomical MRI

High-resolution T2-weighted and T1-weighted images were acquired with an isotropic voxel resolution of 0.7mm^3^ (T2-weighted 3D T2-SPACE scan: TE/TR=565/3200ms; T1-weighted 3D MPRAGE: TE/TR/TI = 2.14/2400/1000ms).

### MRI Preprocessing

#### Neonates

The dHCP data was preprocessed using the dHCP minimal preprocessing pipelines (Makropoulos et al., 2018). Anatomical MRI preprocessing included bias correction, brain extraction using BET from FSL (FMRIB Software Library) and segmentation of the T2w volume using their DRAW-EM algorithm (Makropoulos et al., 2014). The resulted gray and white matter segmentations were used as anatomical masks in further analyses; these masks were manually checked for accuracy.

Minimal preprocessing for the resting-state fMRI included (Fitzgibbon et al., 2016) distortion correction, motion correction, 2-stage registration of the MB-EPI functional image to the T2 structural image, temporal high-pass filtering (150s high-pass cutoff), and ICA denoising using FSL’s FIX (Salimi-Khorshidi, et al., 2014). In addition to this minimal preprocessing, we smoothed the data (Gaussian filter, FWHM = 3mm) across the gray matter, and applied a band-pass filter at 0.009-0.08 Hz. To further denoise the data, we used aCompCor (Behzadi et al., 2007) to regress out physiological noise (heartbeat, respiration, etc.) from the white matter and cerebrospinal fluid (CSF).

#### Adults

HCP data was preprocessed using the HCP minimal preprocessing pipelines (Glasser et al., 2013). For the anatomical data, a Pre-FreeSurfer pipeline was applied to correct gradient distortion, produce an undistorted “native”structural volume space for each adult participant by ACPC registration (hereafter referred to as “acpc space”), extract the brain, perform a bias field correction, and register the T2-weighted image to the T1-weighted image. Additionally, each participant’s brain was aligned to a common MNI152 template brain (with 0.7mm isotropic resolution). Then, the FreeSurfer pipeline (based on FreeSurfer 5.3.0-HCP) was performed with a number of enhancements specifically designed to capitalize on HCP data (Glasser et al., 2013). The goal of this pipeline was to segment the volume into predefined structures, to reconstruct the white and pial cortical surfaces, and to perform FreeSurfer’s standard folding-based surface registration to their surface atlas (fsaverage).

For the resting-state fMRI data, minimal functional analysis pipelines included: removing spatial distortions, motion correction, registering the fMRI data to structural and MNI152 templates, reducing the bias field, normalizing the 4D image to a global mean, and masking the data with the final brain mask. After completing these steps, the data were further denoised using the ICA-FIX method (Salimi-Khorshidi, et al., 2014). To mirror the adult and neonatal preprocessing pipelines, we unwarped the data from MNI152 to acpc space, allowing both groups to be analyzed in “native” space. We then applied spatial smoothing (Gaussian filter, FWHM = 3mm) within the gray matter, band-pass filtered at 0.009-0.08 Hz and implemented aCompCor to regress out physiological noise, just as we did with the neonates.

All subsequent analyses in neonates and adults were performed in each subject’s native space, except for the whole-brain voxelwise analysis.

### Connectivity analyses

We used the 7-network cortical parcellation identified by Yeo et al. (2011). For the whole-hippocampus and long-axis analyses, the hippocampal label was binarized from FreeSurfer’s (surfer.nmr.mgh.harvard.edu) aparc+aseg parcellation and visually inspected for accuracy in each subject. For the first long-axis gradient analysis this label was further sectioned into anterior and posterior portions via manual segmentation using FreeSurfer, with the uncal apex as the dividing marker (Poppenk & Moscovitch., 2011). All labels (cortical networks, hippocampal labels) were originally in CVS average-35 MNI152 space and then registered to each individual subject’s anatomical data using ANTs (Advanced Normalization Tool) 3dWarpMultiTransform (ANTs version 2.1.0; http://stnava.github.io/ANTs; Avants et al., 2011). ANTs is routinely used for developmental dataset registrations (Alexander et al., 2019; Dean et al., 2018). The resulting registrations were checked for accuracy. Similarly, for the long-axis gradient analysis, the hippocampal label in CVS was split into nine equally spaced “slices” along the anterior-posterior axis. Using the same ANTs registration technique for all ROIs provided an extra measure of consistency between groups and between analyses; however, as an added quality check we ran our whole-hippocampus to network analysis using the binarized hippocampal label provided by the dHCP and HCP for each individual. These second results are nearly identical (see Extended Data, Figure 2-1) to the first (Figure 2) thus increasing confidence that our results are not due to registration error.

**Figure 1:**
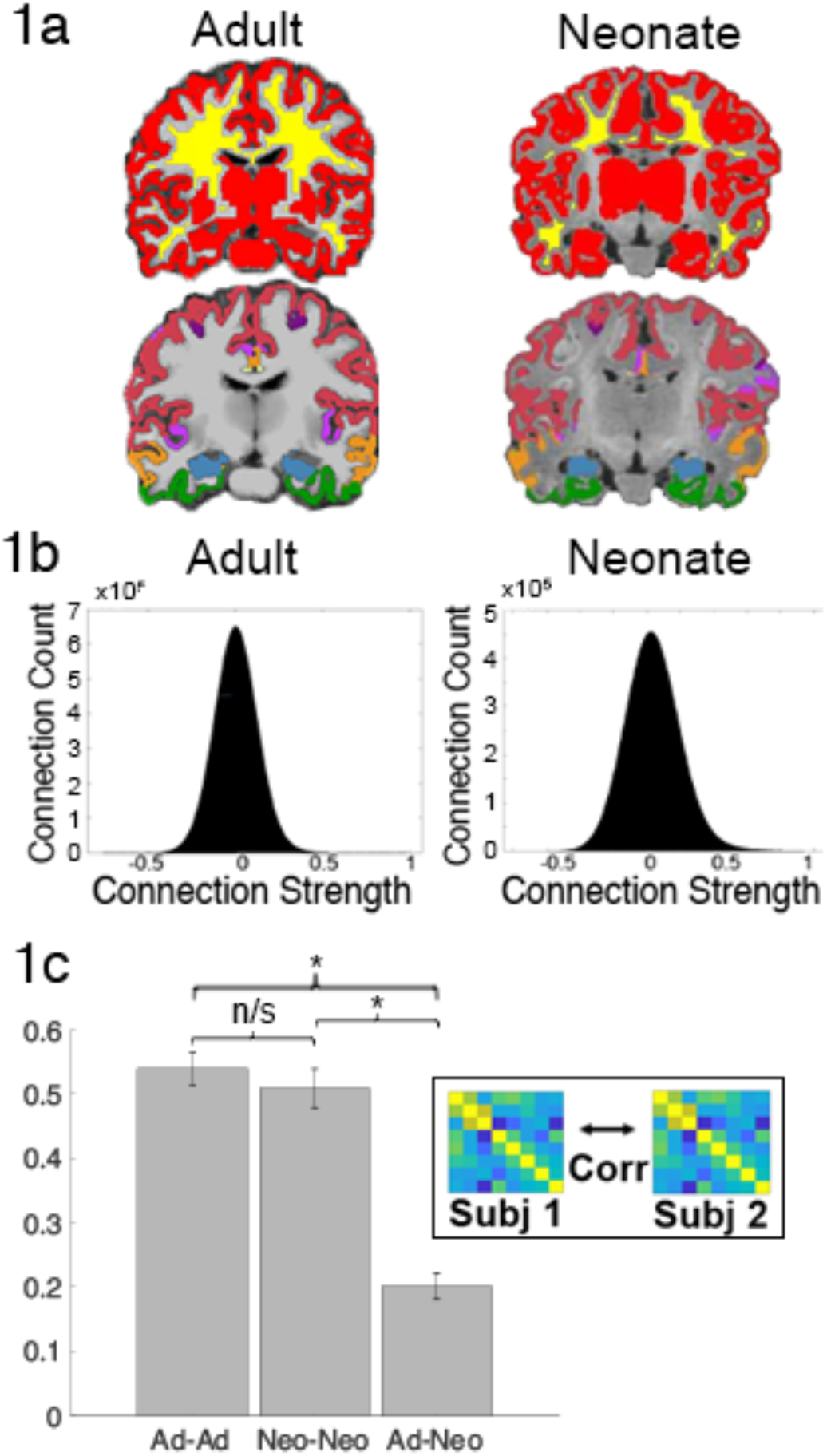
Preliminary Data Checks. a) Gray matter (red), white matter (yellow) and network registrations on the anatomical images of a representative adult and neonate subject; registration image: blue=hippocampus, white = Vis, red = SM, purple (dark) = DA, pink = VA, green = Lim, yellow = FP, orange = DM b) voxelwise correlations distributions of a representative adult and neonate c) between-subject and between-group correlations demonstrate high within-group reliability of connectivity but low between-group reliability between adults and neonates. (*) indicates significance at p<0.05; ns denotes non-significance

**Figure 2:**
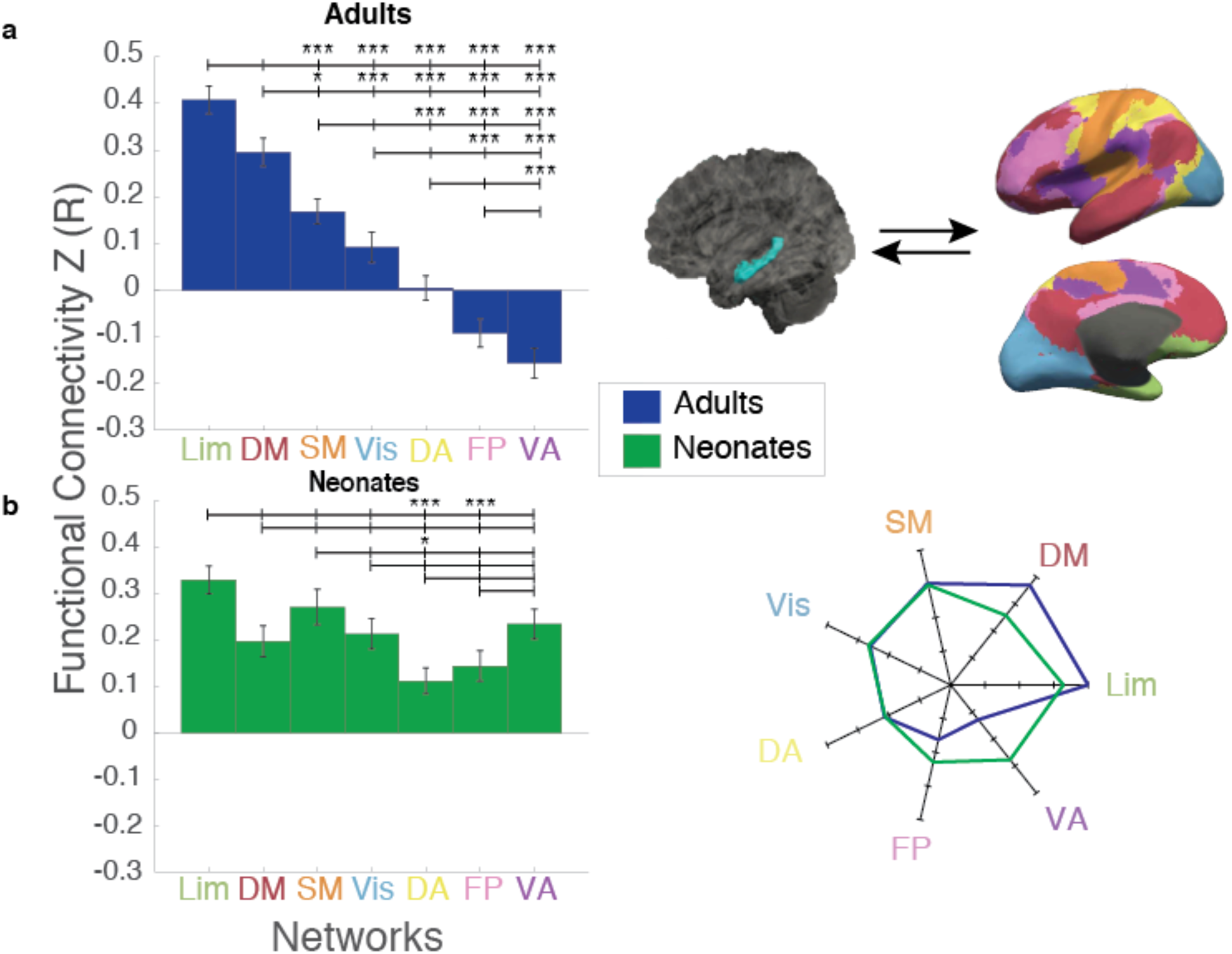
Hippocampal Connectivity to Cortical Networks. a) Comparison of hippocampal connectivity to the seven cortical networks in adults showed a hierarchy of hippocampal connectivity, whereby the highest FC was with Lim, followed by DM, SM, and Vis, almost no FC with DA, and negative FC with FP and VA. b) In contrast, neonates show the same level of FC to almost all of the 7 networks. Rose plot to the right shows adult connectivity compared to neonates to highlight the differences between groups in the pattern of hippocampal FC to these networks. Brain images on the top right depict connectivity between the hippocampus (left) and the seven cortical networks (right). (*) indicates significance at p_HB_<0.05; (***) indicates significance at p_HB_<0.005. Lim=Limbic; DM=Default Mode; SM=Somatomotor; Vis=visual; DA=Dorsal Attention; FP=FrontoParietal; VA=Ventral Attention

After registration to the anatomical data, we registered the labels onto the functional data in neonates using an inverse warp of the func2anat matrix provided by the dHCP. In adults, the labels in acpc space after ANTs registration were then resampled to 2mm cubic voxels to align with the functional data. We manually checked individuals from each sample to ensure the accuracy and fit of the labels to the individual functional data. We extracted the BOLD activation in each label over the time course, averaged within each label, and correlated the hippocampal activity—first whole hippocampus, then along the long-axis (for both anterior-posterior and gradient slices)—with activity in each of the 7 networks to create a Fisher’s Z-scored correlation matrix using Matlab 2018b (The MathWorks, Inc., Natick, Massachusetts, United States).

We also explored differences in the hippocampal connectivity to the whole cortex at a voxelwise scale between adults and neonates to determine whether specific regions within the networks were driving adult-neonate differences. Hippocampal connectivity to the cortex was calculated by correlating the average hippocampal signal and the signal of each voxel within the cortical gray matter mask during the time course for each individual in functional space. To compare the connectivity between adults and neonates, images from both groups were registered to the template space (i.e., CVS average-35 MNI152) before running a between-group analysis. Although this is the only template-space analysis we performed, template-space analyses have been routinely performed to compare infants to adults using similar registration methods (e.g. Gao et al., 2009; Gao et al., 2015a).

### Experimental Design and Statistical Analyses

Where t-tests were performed between regions we corrected for multiple comparisons using the Holm-Bonferroni correction (Holm, 1979); all connectivity values were Fisher’s Z transformed (Fisher, 1915) to normalize the data.

Before doing any of the planned analyses, we first performed data quality checks. To make sure there was no significant motion difference between groups, we calculated the framewise displacement (FD) (Power et al., 2012) based on the six motion parameters estimated from a rigid-body transformation provided by dHCP and HCP. We manually checked the registration of the gray and white matter masks as well as the network and hippocampal labels in the adults and neonates to the registration was accurate. Because we are performing comparisons of correlations between groups, we next wanted to ensure that the correlation distributions were similar and were normally distributed in both neonates and adults; we did this by assessing the correlation of each voxel to every other voxel in the brain and plotting the distribution of those correlations. We also performed between-subject reliability of correlation matrices within and across the adult and neonate groups. We calculated the connectivity of each region (i.e. each of the seven networks and the hippocampus) to every other region for each subject. This connectivity matrix was then correlated with every other subject’s value either between- or within-groups to assess inter-subject reliability; in other words, we correlated the connectivity of every adult to every other adult (within-group) and neonate to neonate, as well as comparing every adult to every neonate (between-group).

Our first analysis examined the relationship of the whole hippocampus to the seven cortical networks. After running a one-way ANOVA with network as the independent variable and connectivity as the dependent variable for both groups, we computed pairwise comparisons between each unique combination of connectivity values to the networks (e.g. Hipp-Lim vs Hipp-VA) to determine networks with significantly different FC to the hippocampus (Snedecor and Cochran, 1989). Rose plots comparing the connectivity pattern of adults and neonates were created by subtracting the mean connectivity across all networks from each individual network (for adults and neonates separately) and plotting the resulting magnitude to show the relative connectivity patterns of the hippocampus to the networks for each group and to compare these patterns between groups.

For our hippocampal-cortical voxelwise analysis, we used FSL’s randomise function to compare between groups and perform permutation testing (to correct for multiple comparisons) in order to determine areas of greater connectivity in adults vs neonates and visa versa. After mapping the individual correlation matrices from subject space into a common template space, we used randomise with default 5000 permutations and clustered the results using FSL’s threshold-free cluster enhancement (TFCE), which corrects for family-wise error (FWE). This produced a list of potential clusters with each cluster’s associated p-value; the p-values were then thresholded at a p< 0.0005, and only those clusters that remained significant after that point are reported in this paper.

For the first long-axis hippocampus analysis, we first computed a two-way ANOVA in each group (separately) using location (i.e. anterior or posterior hippocampus) and network as independent variables and FC as the dependent variable. Pairwise comparisons were then made between the anterior and posterior FC values to each network for each group (e.g. adult antHipp-Lim vs adult postHipp-Lim). For the second long-axis analysis, we conducted a two-way ANOVA at each slice using group and network as independent variables and connectivity as the dependent variable. We also computed a one-way ANOVA at each slice for each group with network as the independent variable. As in the whole-hippocampal analysis, rose plots were created by subtracting out the mean connectivity to all networks (e.g. mean connectivity of adult anterior hippocampus to all networks) from each network and group in the anterior and posterior labels individually to demonstrate comparative connectivity differences between the anterior and posterior regions in each group.

## Results

### Preliminary data-checks

Comparison of the framewise displacement in adults and neonates showed no significant difference of FD between adults and neonates (t(78)=-0.48, p=0.63). Visual inspection of the gray and white matter masks (which are critical for resting-state preprocessing) in Figure 1a shows they are accurately delineating gray/white matter in both neonates and adults; the cortical networks and hippocampal labels also appear to be correctly localized, suggesting that the regions are accurately identified in both neonates and adults (Figure 1a). Figure 1b demonstrates that both neonates and adults have normally-distributed correlation values that are centered around 0. Between-subject reliability of correlation matrices within and across the adult and neonate groups showed the connectivity matrices (i.e. region-to-region connectivity of each of the seven networks and the hippocampus to each other) of each adult subject to each other adult subject were highly correlated, as were the matrices of each neonate subject to each other neonate subject, and a pairwise comparison of subject variability within groups (e.g. adult-adult correlations compared to neonate-neonate correlations) was not significant (t(78)=0.76, p=0.45). But subject-to-subject correlations across the two groups were significantly lower than the within-group correlations (adult-adult vs adult-neo t(78) = 14.09, p=3.87×10^−23^, neonate-neonate vs adult-neonate (t(78)=11.95, p=2.63×10^−19^) suggesting that while the connectivity data are reliable, neonates have different connectivity patterns than adults.

### Whole Hippocampus

We first explored the connectivity of the whole hippocampus to the cortical networks. In adults, there was a main effect of network suggesting that some networks are more strongly connected with the hippocampus than others (Figure 2; one-way ANOVA, F(6,273)=47.11, p=1.84×10^−39^). Subsequent pairwise comparisons showed a clear hierarchy of connectivity, such that hippocampal connectivity was highest to the Limbic (Lim) network (vs hippocampal connectivity to: Ventral Attention or VA (t(78)=12.95, p_HB_=8.42×10^−20^); FrontoParietal or FP (t(78)=11.76, p_HB_=1.20×10^−17^), Dorsal Attention or DA (t(78)=10.09, p_HB_=1.50×10^−14^);Visual or Vis (t(78)=7.20, p_HB_=4.52×10^−9^); and SomatoMotor or SM (t(78)=5.97, p_HB_=7.91×10^−7^)). Hippocampal connectivity to the Default Mode Network (DM) was higher than hippocampal connectivity to: VA (t(78)=10.32, p_HB_=5.80×10^−15^); FP (t(78)=9.07, p_HB_=1.33×10^−12^); DA (t(78)=7.24, p_HB_=4.04×10^−9^); Vis (t(78)=4.63, p_HB_=1.45×10^−4^); and SM (t(78)=3.16, p_HB_=0.014)). Hippocampal-SM connectivity was 3^rd^ highest, and higher than hippocampal connectivity to: VA (t(78)=7.83, p_HB_=3.19×10^−10^); FP (t(78)=6.46, p_HB_=1.09×10^−7^); and DA (t(78)=4.35, p_HB_=3.61×10^−4^)). Hippocampal-Vis connectivity was the next highest (vs VA (t(78)=5.49, p_HB_=5.31×10^−6^); FP (t(78)=4.16, p_HB_=6.38×10^−4^), and connectivity with DA was higher than with VA (t(78)=3.89, p_HB_=1.47×10^−3^). In summary, hippocampal connectivity was highest to Lim, followed by DM, then SM, Vis, and DA; hippocampal connectivity was lowest (i.e. negatively correlated) with the FP and VA networks. In previous literature, the hippocampus is occasionally included as a part of the DM network; our finding of high Hippocampal-DM correlation and anti-correlation between the hippocampus and attention (i.e. FP and VA) networks falls in line with earlier work on the connectivity of the DM network (e.g. Buckner, Andrews-Hanna & Schacter, 2008) and is a good sign of the reliability of our results.

In contrast to the adult pattern, although neonates did show a main effect of network (F(6,273)=5.12, p=2.27×10^−5^), pairwise comparisons indicated that only the Lim and SM networks significantly differ from the rest, with significantly greater connectivity from the hippocampus to Lim vs DA ((t(78)=5.31, p_HB_=2.15×10^−5^) and Lim vs FP (t(78)=4.22, p_HB_=1.33×10^−3^) and significantly greater connectivity to SM vs DA(t(78)=3.35, p_HB_=0.023).

Pairwise comparisons between adults and neonates showed significant differences between the groups, with significantly less connectivity in adults to Vis (t(78)=-2.64, p_HB_=0.040), DA(t(78) =-2.77, p_HB_=0.035), FP(t(78)=-5.33, p_HB_=5.49×10^−6^) and VA (t(78)=-8.62, p_HB_=5.86×10^−13^) networks compared to neonates.

### Hippocampus to Cortex voxelwise analysis

We next explored the connectivity of the hippocampus to the entire cortex at a voxelwise scale; because our previous analysis only focused on 7 canonical networks, we may have missed differences between neonates and adults at a finer grain than that seen on a network level. Thresholding the unpaired t-test results of the whole-brain clusters at p<0.0005 produced 26 significant FWE-corrected (Smith & Nichols, 2009) clusters in the neonates > adults comparison (i.e. 26 clusters where neonatal hippocampal FC significantly exceeds adult hippocampal FC) and 14 significant clusters in the adults > neonates comparison (Figure 3). Specifically, neonates show greater hippocampal FC to frontal and parietal areas, bilateral lingual and pericalcarine cortex and cuneus when compared to adults; frontoparietal differences were particularly prevalent within the right hemisphere. Adults, on the other hand, displayed greater hippocampal FC than the neonates primarily to bilateral isthmus cingulate and precuneus. Cluster sizes and indices for clusters greater than 200 voxels along with peak voxel location and associated brain regions are reported in Figures 3-1 and 3-2 and largely follow the results from the 7-network analysis—the neonatal hippocampus shows greater FC to frontoparietal and attention-relevant areas, whereas the adult hippocampus shows greater FC with regions associated with the default mode and limbic networks.

**Figure 3:**
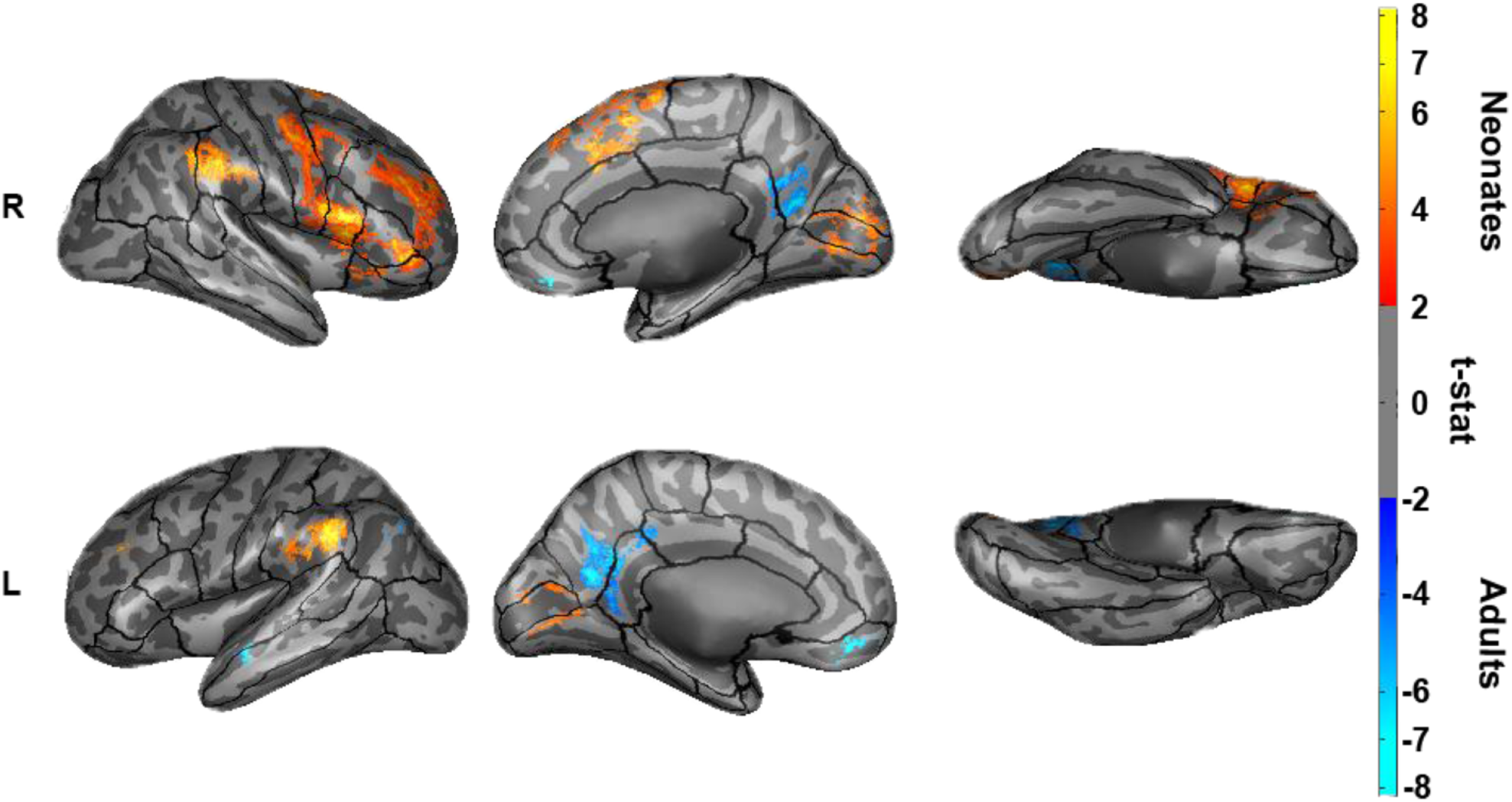
Hippocampal Connectivity to Cortex. Comparison of adult and neonate hippocampal connectivity to the cortex at a voxelwise grain. FWE-corrected results for the contrast of neonate > adult connectivity is shown in warm colors and the contrast of adult > neonate is denoted by cool colors.

### Anterior-Posterior Hippocampus

We next explored the anterior vs. posterior hippocampal connectivity patterns in neonates and adults; previous literature in both humans and other animals suggest functional differentiation of the anterior and posterior hippocampal segments, and thus we may expect these segments to have differences in FC to the 7 cortical networks. In adults, a two-way ANOVA indicated a main effect of network (F(6,546)=60.04., p=5.04×10^−57^) and an interaction between network and anterior/poster hippocampus (F(6,546)=14.31, p=3.54×10^−15^) (Figure 4). In neonates, the two-way ANOVA showed only a significant main effect for network (f(6,546)=7.67, p=6.30×10^−8^) (Figure 4). Pairwise comparisons between the anterior and posterior portions of the hippocampus in adults show greater anterior vs posterior connectivity to the Lim (t(78)=3.53, p_HB_=0.0035), DMN (t(78)=2.38, p_HB_=0.03) and SM (t(78)=3.19, p_HB_=0.0082) networks, and decreased anterior vs posterior connectivity to the DA (t(78)=-3.07, p_HB_=0.0087), Frontoparietal (t(78)=-5.79, p_HB_=9.99×10^−7^) and VA (t(78)=-3.92, p_HB_=0.0011) networks. These results suggest the anterior hippocampus was primarily driving the negative correlations with VA & FP seen at the level of the whole hippocampus in adults. Neonates, however, show no significant differences between the anterior and posterior portions of the hippocampus to any of the networks, suggesting no differentiation/specialization of the hippocampal segments in their connections to the rest of the brain.

**Figure 4:**
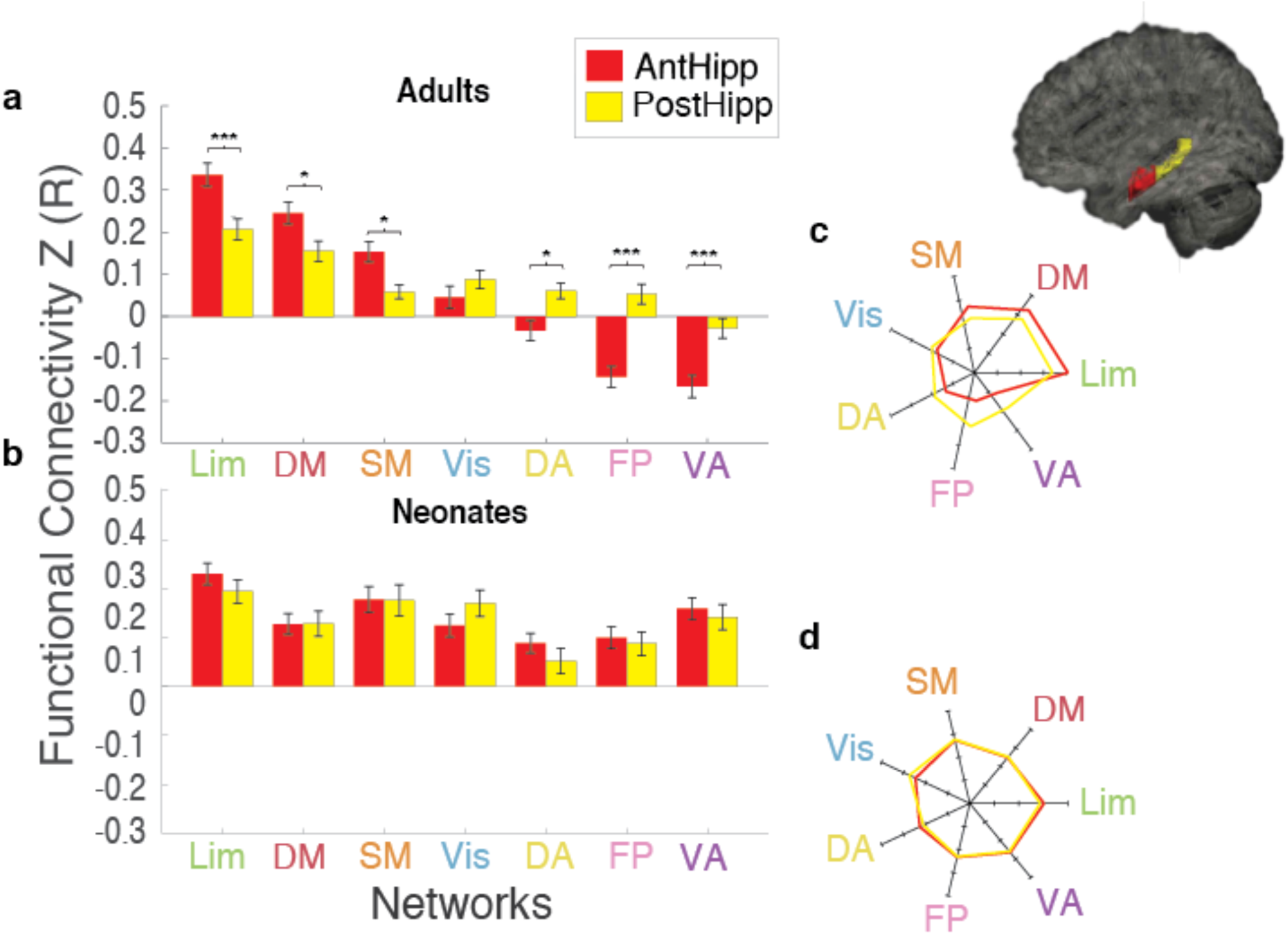
Anterior/Posterior Hippocampal Connectivity to Networks. a) Anterior vs posterior hippocampal-network connectivity in adults b) Anterior vs posterior hippocampal-network connectivity in neonates c) Rose plot comparing anterior vs posterior hippocampal-network connectivity pattern in adults d) Rose plot comparing anterior vs posterior hippocampal-network connectivity pattern in neonates. Brain image to the right shows the anterior (red) and posterior (yellow) hippocampal labels (*) indicates significance at p_HB_<0.05; (***) indicates significance at p_HB_<0.005.

### Long-Axis Gradient

Finally, we investigated the long-axis gradient, which has been demonstrated to map onto a differential functional gradient of the hippocampus. We broke up the hippocampus in each subject into 9 different segments along the anterior-posterior axis and compared the 7-network connectivity to these segments in neonates and adults. Adults showed clear differentiation of network connectivity along the long-axis while neonates showed no clear differentiation (Figure 5). The Lim and DM in adults appeared to have an initial rise and fall of FC along the anterior-posterior gradient of the hippocampus which differentiated them from the Vis, SM, and DA, and the FP and VA showed a similar rise and fall of negative FC along the gradient. One-way ANOVAs for adults and neonates at each slice indicated a main effect of network in adults in all but the most posterior slice (Slice 1 (F(6,273)=23.61, p=1.89×10^−22^), Slice 2 (F(6,273)=40.45, p=4.19×10^−35^), Slice 3 (F(6,273)=50.09, p=2.61×10^−41^), Slice 4 (F(6,273)=49.56, p=5.48×10^−41^), Slice 5 (F(6,273)=49.07, p=1.11×10^−40^), Slice 6 (F(6,273)=25.10, p=1.12×10^−23^), Slice 7 (F(6,273)=13.40, p=2.57×10^−13^), Slice 8 (F(6,273)=5.51, p=2.12×10^−5^)). In the neonates, there was no main effect of network in any of the slices (at p<0.001). To compare between the two groups, we performed two-way ANOVAs (with network and group as independent variables and FC value as the dependent variable) for each of the 9 slices. There was a significant interaction between network and group for the anterior 7 slices (Slice 1 (F(6,546)=7.30, p=1.64×10^−7^), Slice 2 (F(6,546)=16.25, p_=_2.95×10^−17^), Slice 3 (F(6,546)=18.98, p=3.99×10^−20^), Slice 4 (F(6,546)=17.22, p=2.83×10^−18^), Slice 5 (F(6,546)=17.93, p=5.05×10^−19^), Slice 6 (F(6,546)=5.79, p=7.26×10^−6^), Slice 7 (F(6,546)=4.87, p=7.27×10^−5^) and Slice 8(F(6,546)=3.89, p=8.26×10^−4^), but no group differences for the most posterior slice. These results show that the biggest differentiation of hippocampal connectivity to the 7 networks occurs in the anterior 2/3s of the hippocampus in adults and that neonates do not show this differentiation.

**Figure 5:**
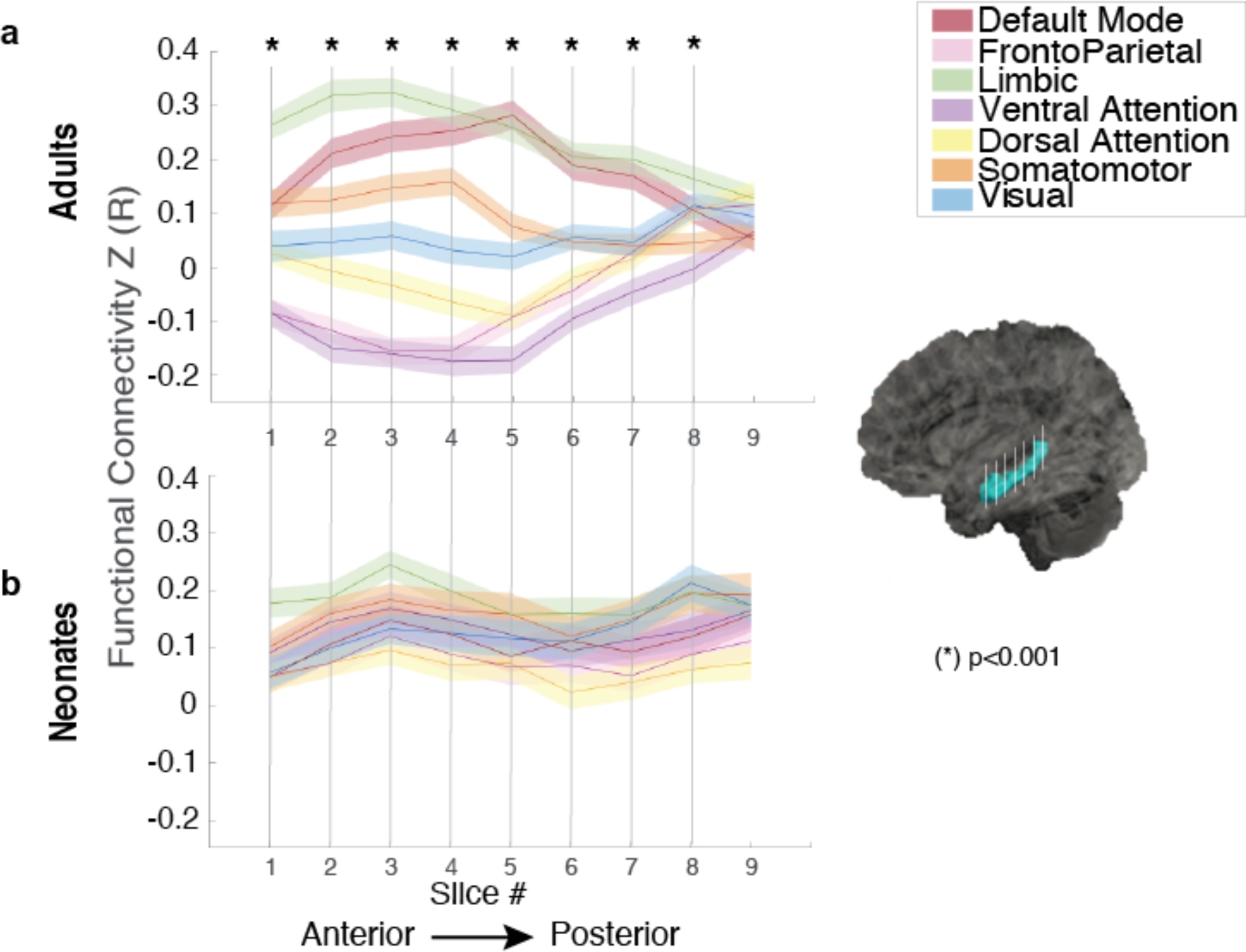
Connectivity along the Long-Axis Gradient to Networks. Comparison of the connectivity of the long axis gradient of the hippocampus to the 7 networks in a) adults and b) neonates. The slices are arranged anterior-to-posterior. Lighter coloring surrounding each line represents the standard error. Brain image on the right demonstrates the hippocampus (blue) segmented into slices (white lines). (*) are slices where the ANOVA shows an interaction between network and group at p<0.001.

## Discussion

Our results show that the intrinsic connectivity of the hippocampal network is not fully mature at birth. Previous functional and volumetric evidence in both non-human primates and humans suggests that the hippocampus continues to develop beyond one year of age, even into middle childhood (Blankenship et al., 2017; Jabes et al., 2011; Keresztes et al., 2018; Lavenex and Banta Lavenex, 2013; Riggins et al., 2016). Although there is some evidence to suggest that the hippocampus is playing a key role in memory formation even early on in rodents (Alberini & Travaglia, 2017; Travaglia et al., 2018), it has been suggested that the long-lasting memories of very young children may be created in a fundamentally different way from adult long-term memories and may rely on cortical mechanisms rather than the traditional hippocampal method (Ellis & Turke-Browne, 2018; Gómez & Edgkin, 2016). Interestingly, multiple studies comparing preterm to term infants show no differences in gray matter volume in the hippocampus with decreased gestational age, implying the better part of hippocampal growth is accomplished prior to birth (Alexander et al., 2019; Ge et al., 2015; Thompson et al., 2008); our work suggests that although the physical bulk of the hippocampus may exist at birth, its connections do not. Specifically, the hippocampus does not have preferential connectivity to any particular network at birth and lacks any long-axis gradient of connectivity, suggesting that the hippocampus, the cortical networks it interacts with, or some combination of both, are immature at birth and may therefore be unable to form long-term memories using adult-like mechanisms. Indeed, the cortex itself is still maturing early on (e.g. Gao et al., 2015b; Ofen et al., 2007; Salzwedel et al., 2019) and it is likely this cortical immaturity, in addition to hippocampal immaturity, is contributing to the differences in memory formation between adults and neonates.

Adults showed a clear hierarchy of FC to the seven networks (consistent with Wael et al. (2018)), whereas neonates lacked this hierarchy. Further, the comparison between adults and neonates shows significant differences between groups to all networks except the SM network, and only marginally significant differences between groups in the Vis and DA networks. The similarity between adults and neonates in connectivity to the SM and Vis networks may be due to the relative maturity of these areas at birth (Arcaro & Livingstone, 2017; Deen et al., 2017; Hurk et al., 2017; Gao et al. 2017, Dall’Orso et al., 2018).

To more specifically determine which regions in the networks were responsible for the differences seen between adults and neonates, we conducted a voxel-wise cortical analysis. Our results indicate that neonates have higher connectivity to much of the cortex as compared to adults with the exception of areas of bilateral medial orbitofrontal, isthmus cingulate and precuneus. This is consistent with Riggins et al.’s (2016) conclusion that 4-year old children rely more on regions “outside” the canonical hippocampal network to complete episodic memory tasks, and other research suggesting the infant cortex is more broadly tuned than in adults (Ellis &Turk-Browne, 2018). The few regions where adults display higher FC than neonates reside mainly within DM network and highlight the immaturity of this network: adults show significantly greater DM-Hippocampal connectivity than neonates, consistent with Gao et al.’s (2015) finding that this network is one of the last to develop in the first year of life.

Our anterior-posterior analysis and long-axis gradient analyses again suggest that the FC differentiation of the hippocampus is lacking at birth. Consistent with previous literature, adults display changes along the long-axis such that the anterior hippocampus shows greater connectivity to the Lim network than the posterior hippocampus but greater posterior vs anterior FC to the attention (i.e. FP and VA) networks; in fact, the anterior hippocampus is especially anti-correlated with these networks, as is consistent with previous literature (Buckner et al., 2009; Wael et al., 2018). The greatest differentiation in FC to the networks in adults occurred within the anterior two-thirds of the hippocampus. In contrast, neonates showed no specificity to any of the networks along the long-axis or the anterior-posterior analysis. Blankenship et al. (2017), Langnes et al. (2018), and Riggins et al. (2016) show evidence of specialization along the longitudinal axis in 4- and 6-year old children but no such evidence is seen in our results, suggesting that maturational changes within the hippocampus may occur before age 4 to produce the preferential connectivity seen in children and adults. Future studies of infants and toddlers can better elucidate when after birth this change in specialization of the long-axis occurs.

Several limitations warrant discussion. A major problem in imaging children is motion artifact. We used the motion-corrected data that were released by the dHCP, took steps in preprocessing to ensure that physiological artifacts were removed from the data in both neonates and adults (Power et al., 2014; Yan et al., 2013), and further motion-matched the neonatal and adult groups. Given that motion-related artifacts are a major confound in FC analyses (Power et al., 2012; Satterthwaite et al., 2013), our approach should minimize the risk of spurious correlations. Other steps we took to minimize potential confounds included visual inspection of spatial registration results (and using established registration procedures that have been previously performed on infants (Alexander et al., 2019; Dean et al., 2018; Gao et al., 2009; Gao et al., 2015a)), performing the analyses in the native-space of each individual, and checking the reliability of the correlation values across participants in each group to ensure they were not particularly noisy in the neonatal group. A result of particular note is that neonates showed primarily positive FC from the hippocampus to the networks, while adults showed slightly negative FC for some networks. Blankenship et al., (2017) similarly fail to show any negative hippocampal FC in their sample of 4- and 6-year old children (but this may be due to their preprocessing steps, see Murphy & Fox, 2017 for discussion). Here, we used the same preprocessing steps in both neonates and adults and used aCompCor and other preprocessing steps that should not necessarily remove negative correlations if they were there. Indeed, we found a normal distribution of correlation values in both neonates and adults (Figure 1) suggesting that negative correlations do exist in neonates, but not between the hippocampus and the cortex. Further, regardless of the negative vs. positive correlation differences we observe a difference in the pattern of FC in adults (demonstrated in the rose plots) primarily in the anterior portion of the hippocampus; this is missing in neonates.

Differences in arousal states between the groups present another challenge. Mitra et al., (2017) showed differences in resting-state connectivity between sleeping infants and waking adults. However, observation of Mitra et al.’s data suggests although the magnitude of connectivity may differ between arousal states, the overall pattern of connectivity remains similar (i.e. the same clusters of connectivity are observed in sleep and in rest and their relative comparison to other clusters remains similar across sleep states and age groups). Further, although notable differences are seen between the 24-mo sleeping infants and waking adults in Mitra et al., this difference is far less pronounced in the younger 6-mo infants. Based on previous EEG studies (e.g. Roffwarg, 1966), it is possible that younger infants experience less slow-wave sleep and more REM sleep and thus, younger infants (vs. older infants) during sleep would be expected to look more like awake adults due to the high similarity of REM and wakefulness activity patterns in EEG, particularly in infants. Because we would expect more awake-like REM sleep and less slow-wave sleep in young infants, we believe that the neonates in the current study are unlikely to show major wake/sleep confounds in their connectivity patterns. Finally, analysis of the same dataset but specifically of visual network connectivity showed striking *similarities* in connectivity patterns (https://www.biorxiv.org/content/10.1101/712455v1) and therefore further suggest that any differences in data acquisition and/or sleep states between adults and neonates are unlikely to systematically lead to the differences in network connectivity that we find here.

Finally, we found the motion- and gender-matched HCP adults used this manuscript tended to have lower tSNR than their respective dHCP counterparts. To ascertain this discrepancy was not the cause of observed group differences, we identified a separate group of 40 HCP adults whose tSNR matched that of the 40 neonates used here and performed our whole hippocampus to network analysis on this group. The resulting pattern matches the pattern observed from the previous analyses (i.e. using the original 40 HCP adults), reiterating that identified differences in connectivity patterns between adults and neonates are likely not spurious byproducts of discrepant data quality (Extended Data, Figure 1-1).

In conclusion, our results suggest that the resting-state FC patterns of the human hippocampus are immature at birth. This immaturity may play a key role in infantile amnesia and the vast differences between adults and neonates shown here suggests a fundamentally different memory and learning system from that of adults may be present at this point in development.

## Acknowledgements

The authors declare no competing financial interests

## Legends

**Figure 1-1:**
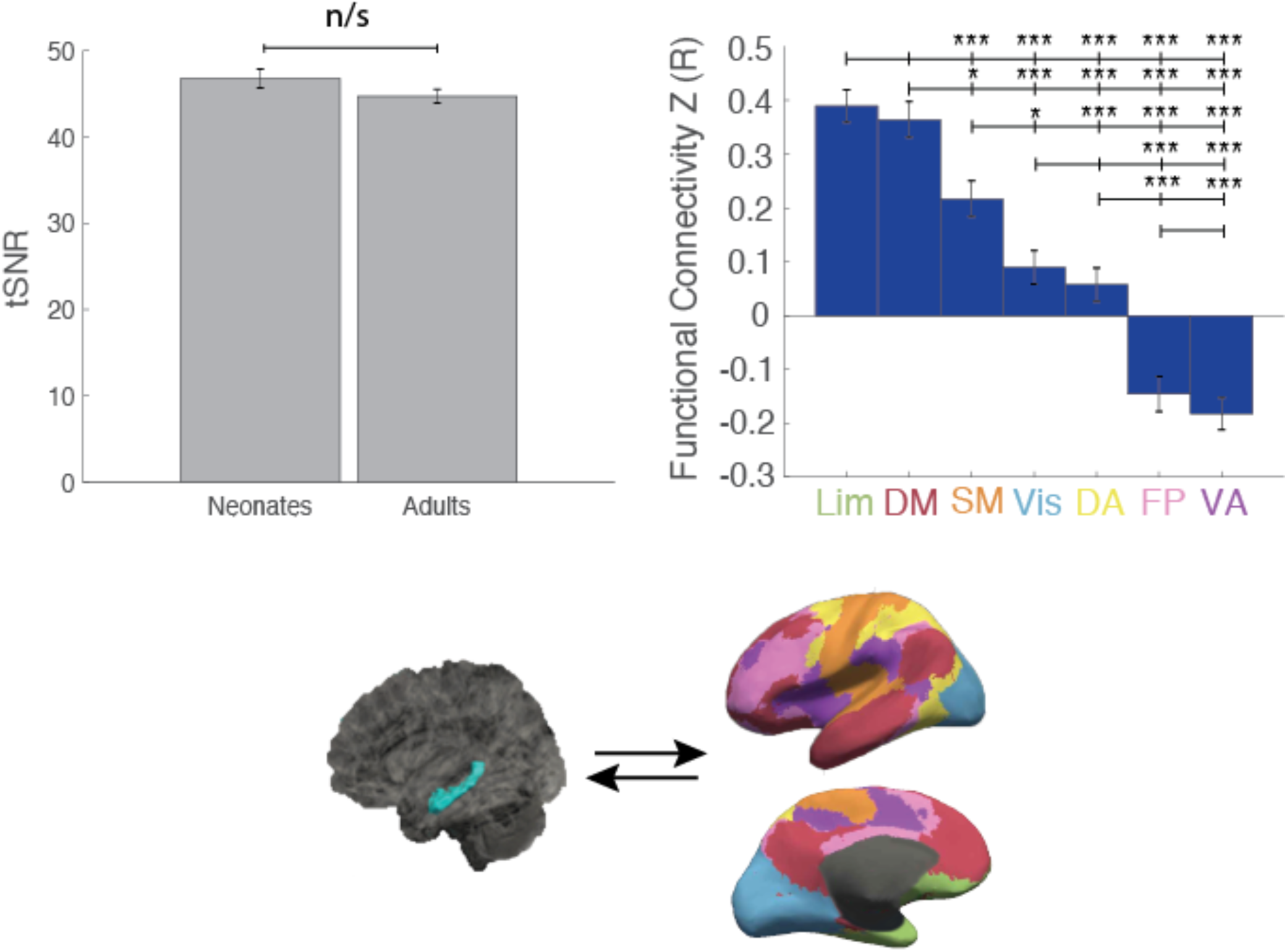
tSNR-Matched Adult Hippocampus to Networks. Hippocampal-network connectivity of 40 tSNR-matched HCP adults again shows very similar results to the motion-matched and binarized-hippocampal analyses. Hippocampal connectivity in adults shows a clear hierarchy, with strong positive connectivity to Lim and DMN and negative connectivity to FP and VA (*) indicates significance at pHB<0.05; (***) indicates significance at pHB<0.005.

**Figure 2-1:**
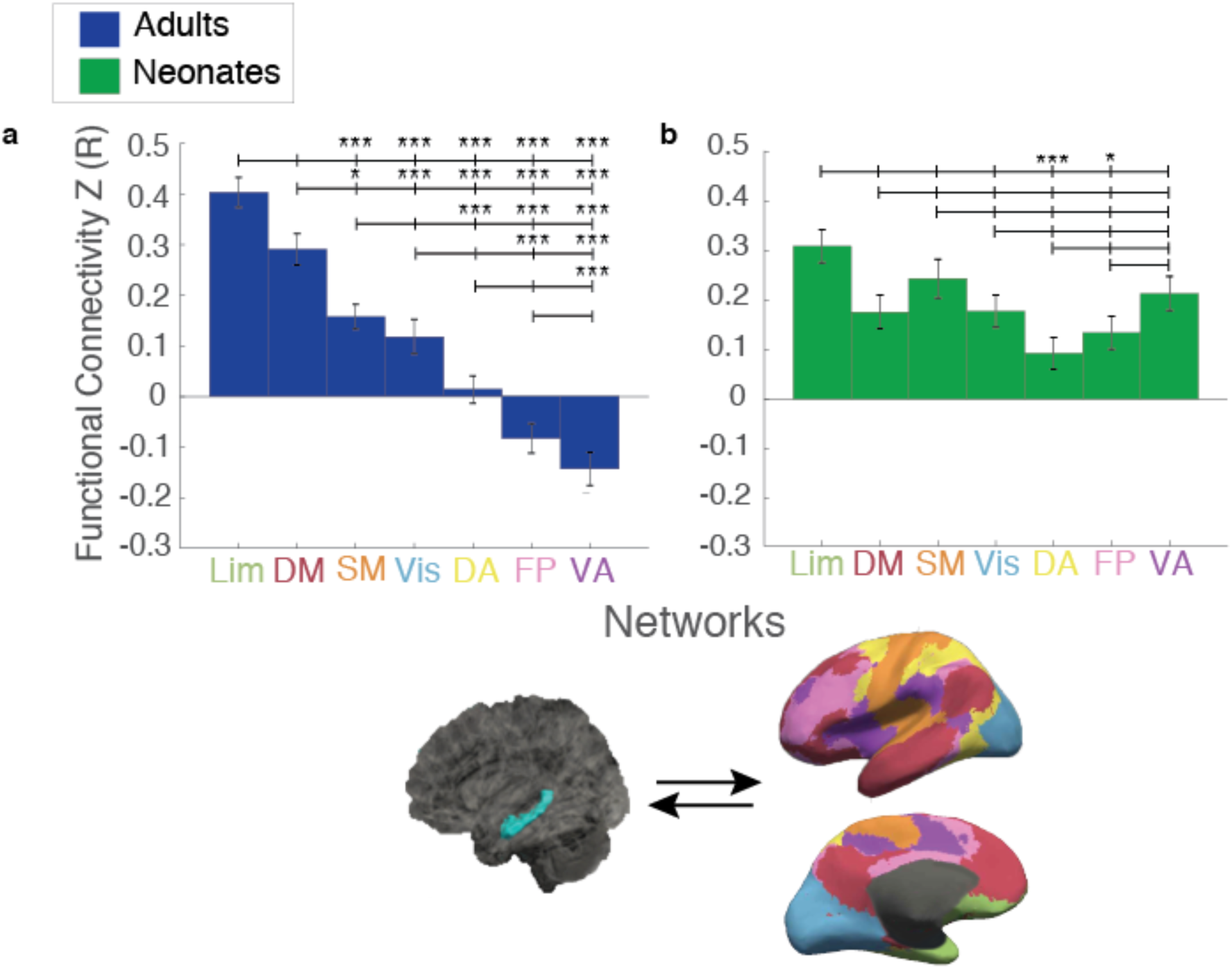
Binarized Whole Hippocampus to Networks. Hippocampal connectivity to the networks using a binarized HCP/dHCP hippocampal ROI yields very similar results to the ANTs registered hippocampal ROI (see figure 2). As with the initial analysis, hippocampal connectivity in adults shows a clear hierarchy whereas neonates display very few differences in hippocampal connectivity strength to the networks. (*) indicates significance at pHB<0.05; (***) indicates significance at pHB<0.005.

**Figure 3-1:**
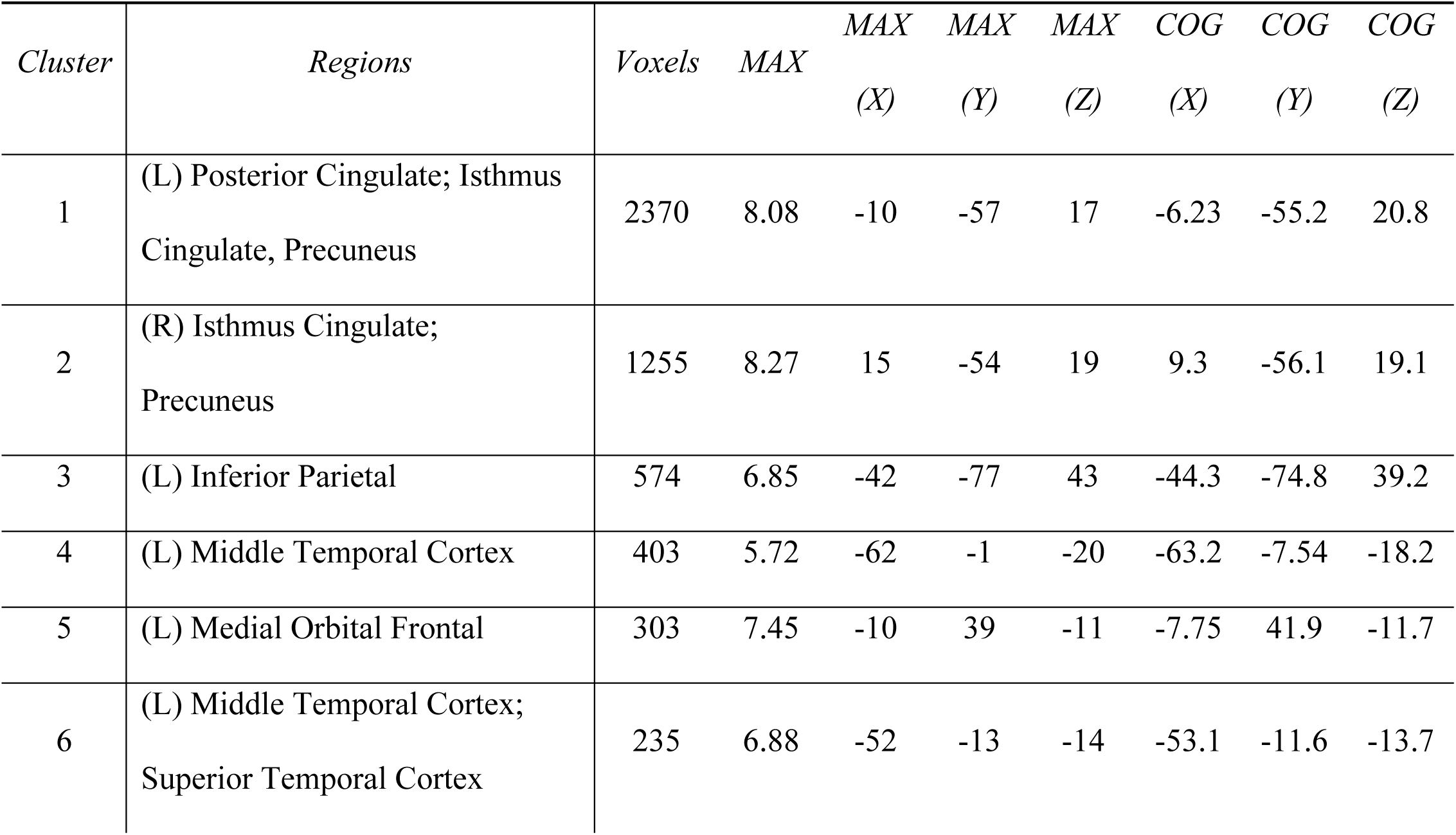
Hippocampal Connectivity to Cortex, Adult>Neo. Clusters are listed from largest to smallest. Peak coordinates (MAX) are listed in MNI space as well as center of gravity (COG) for each cluster

**Figure 3-2:**
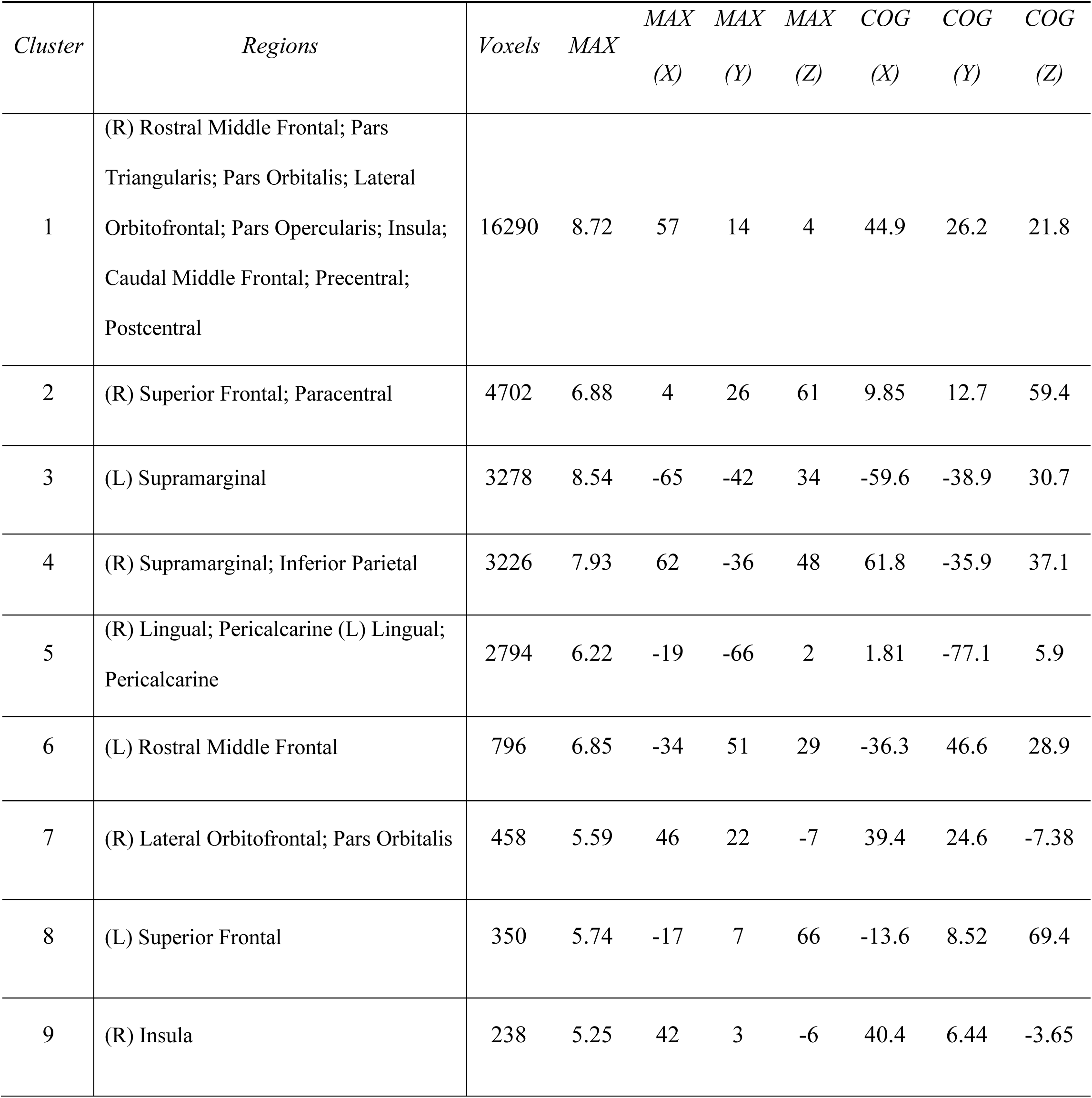
Hippocampal Connectivity to Cortex, Neo>Adult. Clusters are listed from largest to smallest. Peak coordinates (MAX) are listed in MNI space as well as center of gravity (COG) for each cluster

## Notes

### Competing Interest Statement

The authors have declared no competing interest.

### Summary of Updates

Additional data quality checks concerning tSNR differences and hippocampal ROI identification were made; results strengthen additional findings. Additional literature added to the introduction and discussion

